# IntegrAlign: A comprehensive tool for multi-immunofluorescence panel integration through image alignment

**DOI:** 10.1101/2025.04.10.648082

**Authors:** Leo Hermet, Leo Laoubi, Martial Scavino, Anne-Claire Doffin, Alexia Gazeu, Justine Berthet, Berenice Pillat, Stephanie Tissot, Sylvie Rusakiewicz, Marie-Cécile Michallet, Nathalie Bendriss-Vermare, Jenny Valladeau-Guilemond, Jean Hausser, Christophe Caux, Margaux Hubert

## Abstract

**Motivation:** Tyramide-based multiplex-immunofluorescence (mIF) enables the simultaneous analysis of up to seven protein markers on a whole slide, providing a comprehensive approach to study the tumor microenvironment. Integrating multiple mIF panels through image alignment of serial slide significantly expands the number of cell populations analyzed in a single space. IntegrAlign was developed to optimize this integration on serial whole slides, enhancing the value and applicability of mIF for comprehensive spatial analyses and enabling biomarker discovery at scale.

**Results:** IntegrAlign, leveraging the SimpleITK toolkit, applies a two-step alignment using rigid and B-spline transformations to integrate serial mIF whole slides. Validation on simulated and real datasets demonstrated alignment accuracy below the diameter of a cell nucleus (∼6 µm), outperforming existing methods. This precision enhances spatial analyses by combining extended phenotypic data, supporting novel insights into tissue architecture and cellular interactions.

**Availability and Implementation:** IntegrAlign is open-source, implemented in Python, and freely available under the MIT license at https://github.com/CAUXlab/IntegrAlign.

## Introduction

Tissue histopathology, in particular hematoxylin and eosin (H&E) staining and immunohistochemistry (IHC), have long allowed visualizing the microscopic structures of tissues and remain central to the diagnosis of many diseases, especially cancer. Traditionally, tissue images are examined by anatomopathologists to evaluate the overall architecture and identify structural abnormalities. To support this goal, machine learning and artificial intelligence (ML/AI) are being developed to automatically extract spatial and architectural information from tissue images, moving towards a computer-aided diagnosis.

Despite these advances, H&E only visualizes tissue architecture and IHC is limited to small number of markers. To address this, highly multiplexed tissue imaging methods (such as CODEX [1], MIBI [2], IMC [3]) were recently developed to profile many markers and cell populations on a single slide. However, highly multiplexed imaging techniques suffer from high price, long image acquisition time and small size of analyzed areas which require focusing on small regions of interest (ROI), all of which limits their clinical relevance and make them inapplicable to large cohorts of patients. Multiplex-immunofluorescence (mIF) offers an attractive trade-off, allowing to analyze multiple markers and populations (up to 7 markers stained per slide) on a whole slide for large cohorts in a reasonable time frame and cost.

It is classical to perform mIF on serial slides with complementary panels of markers to obtain a more detailed tissue profiling. However, the most common and simplest way to interpret the mIF data is an individual analysis of each panel, which reduces the potential for in-depth spatial analysis. To overcome this limitation, data from different mIF panels performed on serial slides can be integrated. This requires image alignment, also called registration, to process two different data sets into a common coordinate system: registration estimates a transformation which maps points from one image to the corresponding points in another image. This method has previously been applied to consecutive staining on a same slide generating multiple images [4-5] and to biomedical images such as MRI brain scans [6-9]. However, the registration of images coming from serial tissue slides is challenging due to several factors related to tissue preparation, including morphological changes (compression or stretching), missing or damaged areas, as well as tissue rotation and translation. While rigid registration methods are defined as geometric transformations that preserve all distances, non-rigid registrations enable transformations with image deformation. Such strategies have previously been applied to serial H&E slides [10-12], but never using mIF images generated with distinct panels of markers and fluorochromes.

In this article, we introduce IntegrAlign, a novel method for integrating two mIF panels by image alignment using the SimpleITK toolkit [13]. We present the first open-source pipeline developed to augment the analytical capabilities allowed by the mIF technique. Through a comparison with existing algorithms based on manual or automated detection of anchor points, we clearly demonstrate the effectiveness and accuracy of IntegrAlign. By precisely aligning mIF images, integrating multiple panels using IntegrAlign increases the number of cell populations that can be analyzed simultaneously (**Fig. 1A**). IntegrAlign will synergize with a fast and inexpensive spatial analysis of whole tissue slides by mIF to open new avenues for the comprehensive analysis of spatial biology and the identification of new biomarkers in large cohorts of patients.

**Figure 1:**
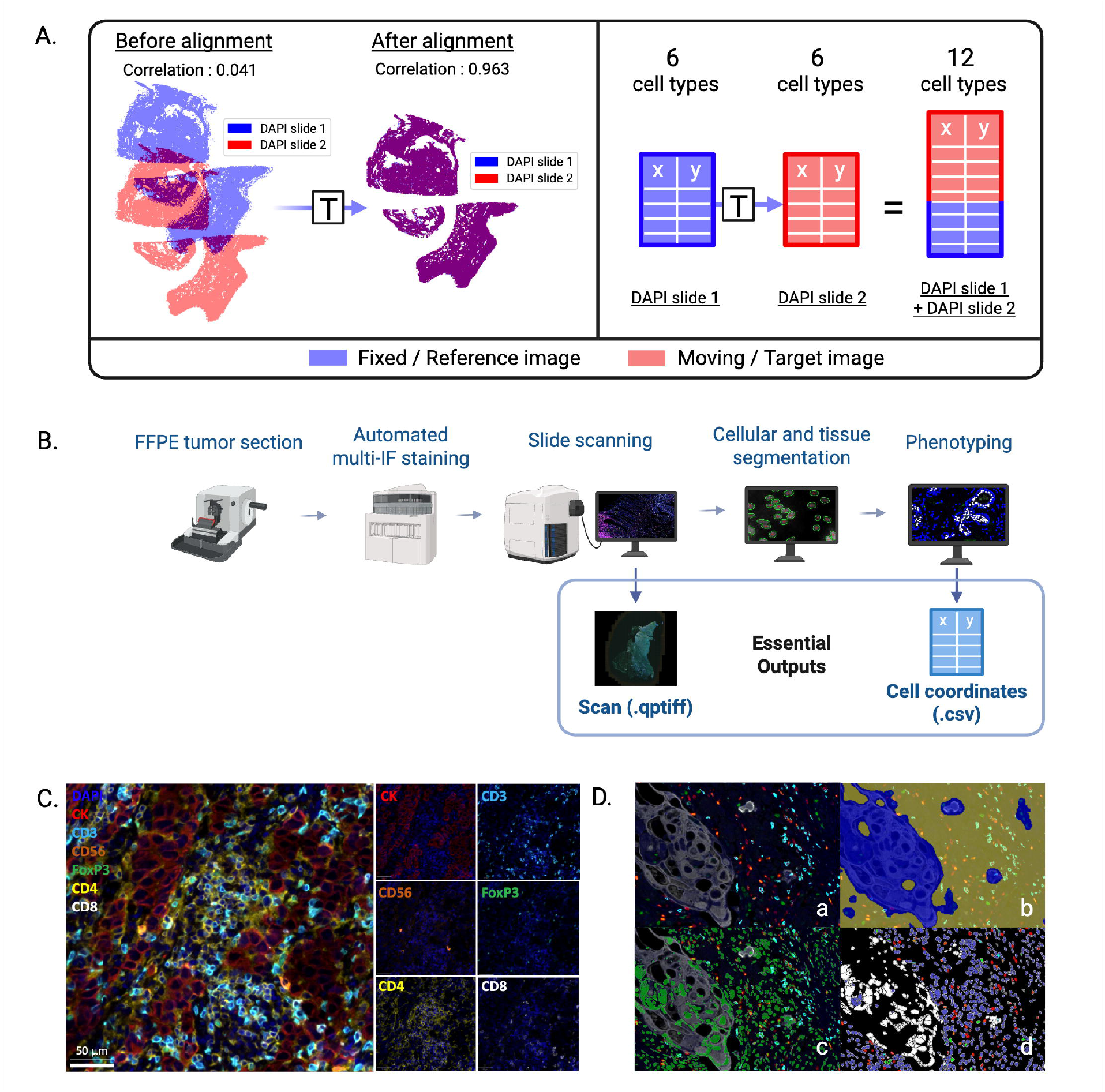
Principles of Integralign tool and multiplex-immunofluorescence analysis. **(A)** Objective of the Integralign tool for multiplex-immunofluorescence analysis. This panel illustrates how Integralign accurately matches cell coordinates (DAPI) between two serial slides, showing the correlation factor between the rasterized cell positions of the two slides before and after alignment. Once alignment is done all cell types are merged, resulting in a total of 12 distinct cell types in the coordinate system of the target slide. **(B)** Workflow of multiplex-immunofluorescence analysis from tissue to spatial data. **(C)** Illustration of a 7-color opal-based multiplex-immunofluorescence staining. **(D)** Representation of tissue segmentation (D.b), nuclei segmentation (D.c) and phenotyping (D.d) from a scan (D.a) in breast tumor.

## Materials and methods

### 1 Image acquisition and data acquisition

In this section we describe the different steps required to generate input files for the IntegrAlign pipeline (**Fig. 1B**). Formalin-fixed paraffin-embedded (FFPE) tumor tissues have been collected at diagnosis from surgical specimens. Written informed consent for the use of samples for research purposes were obtained from all patients prior to analyses. For each tumor (except neuroblastomas), a pair of two 4µm thick serial sections were stained by Opal-based multiplex-immunofluorescence (mIF) to detect cell types corresponding to the associated panel (**Table 1**). To design the mIF panels, specific markers were chosen based on previous literature data [14]. Briefly, the seven-colors mIF assay was conducted with sequential staining cycles on an automatic stainer. Stained FFPE samples were then scanned and analyzed using InForm or IFQuant software (**Fig. 1C-D**): (i) tissue segmentation was first performed to define tissue and empty/artefact areas, (ii) then cell segmentation was achieved using nucleus recognition and expansion, (iii) and finally cell phenotyping was performed to attribute mutually exclusive phenotypes to each cell.

**Table 1.**
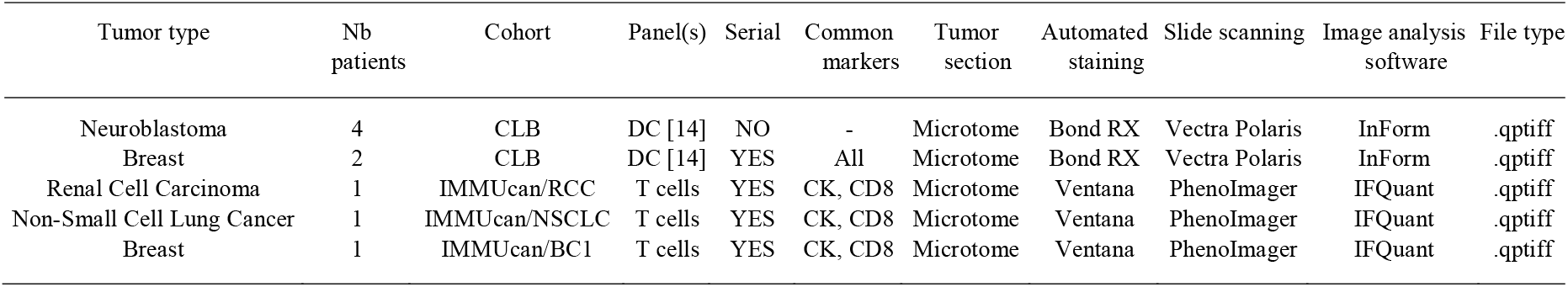
Data overview. Panel DC corresponds to 7markers: DAPI, CLEC9A, BDCA2, CLEC10A, CD8, CK (Cytokeratin), DC-LAMP as already described in Hubert et al. [14]. Panel T corresponds to 7markers with 2 different panel for each slide, (1): PD1, CK, Ki67, GB, CD8, PDL1; (2) CD4, CK, CD3, CD56, CD8, FOXP3

### 2 IntegrAlign pipeline

IntegrAlign pipeline leads to the integration of cell coordinates from 2 serial slides in the same system using scans alignment. **Figure 2** summarizes the different steps of IntegrAlign pipeline that are detailed in the following sections.

**Figure 2:**
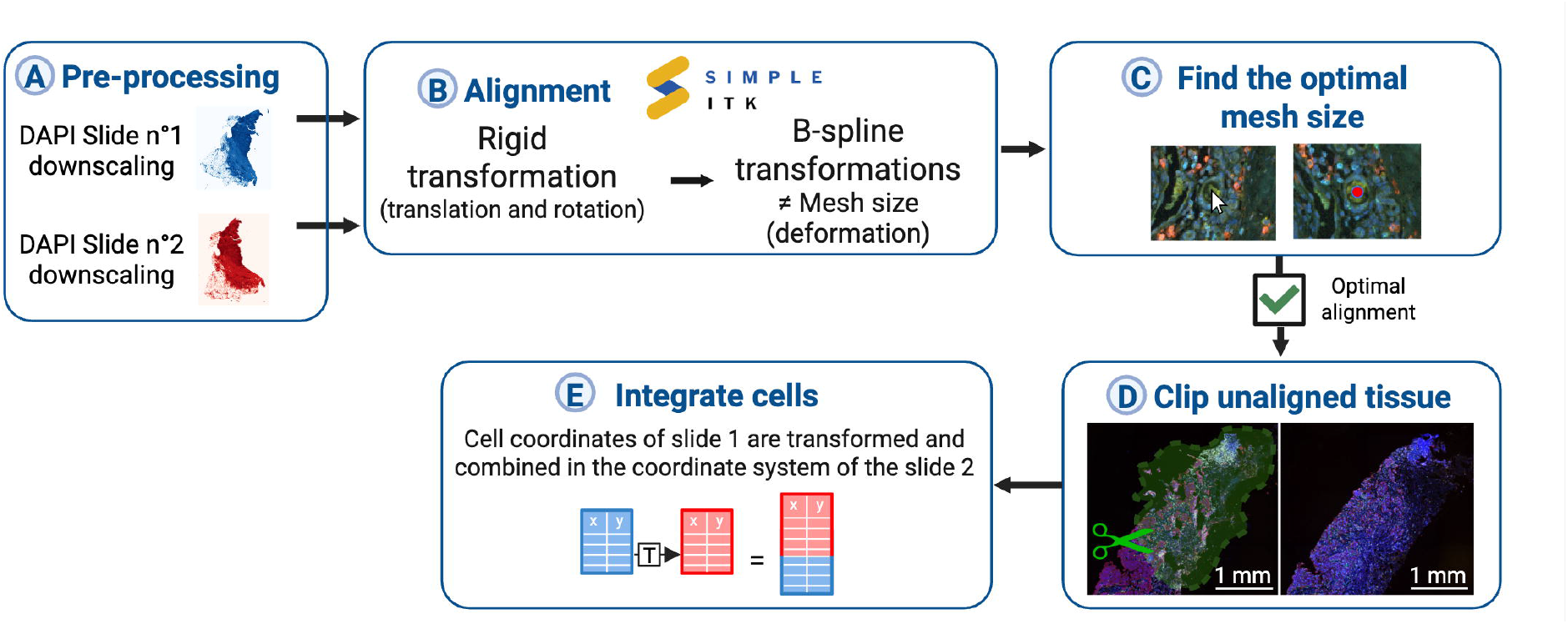
IntegrAlign pipeline. Overview of the IntegrAlign workflow composed of 5 main stages. (A) In the first stage, **DAPI** channel images are downscaled (1/32e) for both serial slides to save computational time and retain only the relevant information. **(B)** Then, downscaled serial slides are aligned using the Rigid and B-spline transformations. **(C)**Several B-spline transformations are evaluated and optimal mesh size is selected based on the quality of deformation and cell-to-cell matching. Indeed, quality control images showing strength and direction of induced deformations are available for visual checking. Alignment accuracy is verified using the napari viewer displaying both slides with the cursor transformed from first to second slide. **(D)** After achieving optimal alignment, using the same dual slide napari viewer, non-alignable tissue sections are excluded by manual clipping. **(E)** Finally, coordinates of both serial slides are merged by transforming those of slide n ° l into slide n ° 2 and removing the ones in excluded areas.

### 2.1 Image processing

To achieve the alignment of two images, it is essential to first reduce the complexity of the images, which are originally composed of 8 high-resolution channels. The DAPI channel, which highlights cell nuclei, was selected because it is the mandatory common marker between serial slides. Also, we chose DAPI among all common markers because of its ubiquitous expression of cellular information. While the full-resolution images exhibit a discrepancy in the spatial arrangement of cells between the two serial slides at high magnification (corresponding cells from serial slides n°1 and 2 failing to match in x60 magnification, **Fig. 3A**), we observed that the cell density patterns remain consistent at lower magnification (x10, **Fig. 3A**). Consequently, to eliminate unnecessary single-cell information while preserving the overall density patterns, we used a down-sampled version of the DAPI channel (1/32^th^ of the original resolution), which was encapsulated within the qptiff-formatted image (contains multiple resolutions of the original image for pyramid representation). This process reduced the image size from 8 channels at 48960×36480 pixels to 1 channel at 1530×1140 pixels, sharply decreasing computational resolution time while preserving critical information required for accurate alignment.

**Figure 3:**
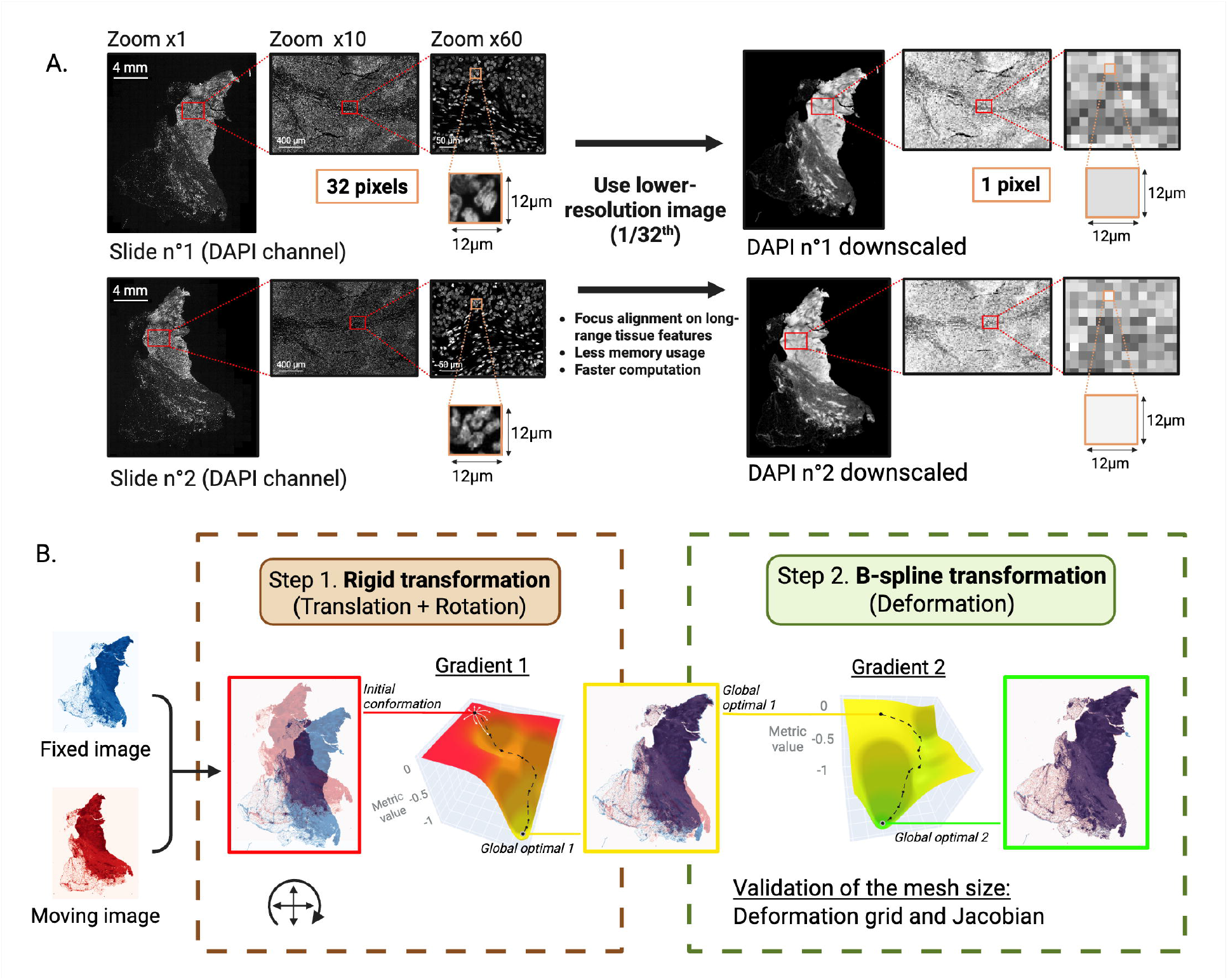
Image pre-processing and alignment with the package SimpleITK. **(A)** DAPI channel images of two serial slides at x l, x l0 and x60 zooms for orginal image (resolution for slide 1 = 48960×36480 pixels) on the left and downscaled image (1132th; resolution for slide 1 downscaled = 1530x l 140 pixels) on the right. Red and orange rectangles indicate the zoomed-in areas before and after downscaling. **(B)** Representation of the alignment of two downscaled serial images using SimpleITK package including two sucessive transformations with an optimisation algorithm employing pixel-wise correlation metric: 1) Euler2DTransform (rigid transformation) performing translation and rotation, followed by 2) deformation using BSplineTransform (B-spline transformation). Visualization of the moving images at global optimal 1 and 2 are from resampling.

### 2.2 Alignment

The goal of alignment is to map points from one image to their corresponding points in another image by estimating the transformation of the coordinates. This can be achieved using the SimpleITK toolkit (https://github.com/SimpleITK/SimpleITK) [13] and in particular three key components: the optimization algorithm, the similarity metric, and the transformation.

#### 2.2.1 Optimization algorithm

In order to define the optimal transformation, we used an optimization algorithm called the Limited-memory Broyden-Fletcher-Goldfarb-Shanno with Bound constraints [15] (L-BFGS-B). It corresponds to a gradient-based optimization algorithm commonly used in numerical optimization that belongs to the class of quasi-Newton methods, designed for optimizing smooth, non-linear functions. The addition of support for simple box constraints on variables in L-BFGS-B algorithm, helps to ensure that the transformation remains within safe and realistic boundaries. From an initial conformation of the fixed and moving images, the L-BFGS-B algorithm efficiently calculates the optimal direction for parameter adjustments, using derivatives (gradient illustrated in **Fig. 3.B**) and an approximation of the Hessian matrix, to maximize the correlation (see similarity metric) between the images at each iteration. The global optimum is defined at convergence of the similarity metric, representing the ideal transformation of the fixed image within the moving image’s coordinate system to maximize the correlation.

#### 2.2.2. Similarity metric

In SimpleITK [13], alignment relies on the values of pixels intensity in both images. This is particularly relevant in our case where the images of serial slides have similar structures and content (DAPI) but differ in terms of pixel intensity. In order to measure how closely two images resemble each other we used the similarity metric called normalized cross-correlation coefficient which quantifies the similarity of pixel intensities between images after normalization. This metric is defined as:

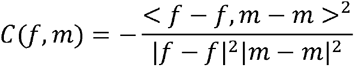

in which, *f* and *m* are the vectors of image pixel intensities. The minus sign makes the metric to optimize towards its minimal value. The correlation is computed efficiently without requiring square root calculations, making it practical for optimization purposes.

This similarity metric is computed between a fraction of the pixels from the fixed and moving images. Consequently, to compute the similarity metric at each iteration, we need the corresponding transformed moving image.

This image is obtained through resampling: A regular grid of points is defined using the pixels of the reference/fixed image. At each iteration, all pixel coordinates from the grid can be transformed (using the corresponding parameters) to the target image with the direct transformation. Then, a linearly interpolated value is computed from the surrounding points in the target image to obtain the corresponding intensity value (pixel’s value) that will be imported back (mapped) to the original coordinate (in the fixed image system). This results in a resampled image that can be used to compute the similarity metric at each iteration between fixed and moving (resampled) image.

#### 2.2.4 Transformation

While the similarity metric quantifies the likeness between images, the optimization algorithm needs also a transformation that involves adjusting parameters to dictate how one image aligns with another. In SimpleITK [13], the transformation is done from the physical virtual space (corresponding to the physical space of the reference image in our case) onto the physical space of the target image.

Here we employ composite transformations, with the sequential application of 2 types of transformation. This approach allows to first implement partial alignment and then introduce more complex deformations. Indeed, the first step was to use a Rigid transformation (**Fig. 3B**) which encompasses rotation and translation adjustments. It can be defined as:

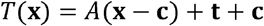

Where *A* is the rotation matrix, **c** the fixed center around which the rotation occurs and **t** the translation vector. This partial alignment step serves as a foundational stage, mitigating initial misalignments between the images. Indeed, applied to the optimization algorithm of image registration, it leads to the global optimal 1 (**Fig. 3.B**) which is the result of the best conformation for correlation between fixed and moving images with a rigid transformation. Then, as a second step, we can start from this optimum to integrate deformations using a B-spline transformation (**Fig. 3B**). Here, the deformation field is modeled using B-splines, allowing local deformation through a grid of control points. Indeed, the deformation is defined by a sparse grid of control points 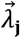, each influencing a specific deformation 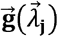. We can regulate the number of control points and thereby modulate the resolution of the transformation by adjusting the mesh size. Using a cubic spline order (degree of the piecewise polynomial) by default, the mesh size parameter corresponds to the number of polynomial patches comprising the finite domain of support and can be defined as:

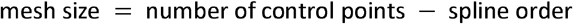

Next, using a B-spline interpolation kernel, we applied the deformations defined by the control points 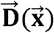 to any point in 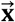 the image.

Using those two transformations subsequently results in the global optimal 2 (**Fig. 3.B**) with a final conformation corresponding to a maximized correlation between fixed and moving images after translation, rotation and deformation.

### 2.3 Definition of the optimal mesh size

For the B-spline transformation we can induce different level of deformation depending on the number of control point using the mesh size parameter. We used different strategies described in this section to determine the most suitable mesh size and to provide information about the overlapping images, the induced deformations, as well as the cell-to-cell matching.

#### 2.3.1 Visualization of overlapping slides

First, for a comprehensive evaluation of the alignment, IntegrAlign allows the visualization of the overlay between both images —fixed and moving— (**Fig. 3B**). DAPI channel downscaled image from slide n°2 was resampled using SimpleITK to retrieve the aligned moving image to the fixed image. Then, the pixels of the resampled image of DAPI from slide n°2 downscaled were combined with the image of DAPI from slide n°1 downscaled using alpha blending.

#### 2.3.2 Deformation grid

To determine the most suitable mesh size, we integrated the visualization of the deformations (illustrated in **Fig. 5.D** left) in the IntegrAlign pipeline with a transformed grid overlaying the resampled moving image. In addition, we showed the control points (blue dots) and their associated displacement (black arrows). This visualization helps identifying areas of deformation and their relevance to the needed adjustment of the slide.

#### 2.3.3. Jacobian

While the deformation grid provides an overview of how the tissue is being deformed, we included in IntegrAlign a more intuitive information about the specific locations of these deformations within the tissue. Indeed, using SimpleITK, we computed the Jacobian determinant. Briefly, from the B-spline transformation we generated the corresponding vector field of displacements from which we determine the gradient (rate of change of displacement). Then, we evaluated the determinant of the resulting matrix, denoted as *Jacobian:*

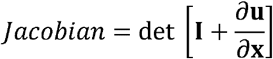

Where **I** the identity matrix and 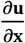 the gradient of the displacement field. This provided a scalar image where each pixel’s value represents the local deformation magnitude (**Fig. 5.D** - right panel). In this case, a determinant greater than 1 indicates compression, signifying that the tissue is being squeezed or compressed at that location. Conversely, a determinant less than 1 corresponds to expansion, indicating that the tissue is being stretched or expanded. This information allows to verify the coherence and accuracy of the deformations, providing insights into the dynamic changes occurring within the tissue.

#### 2.3.4 Duplicated cursor for cell-to-cell matching

Last step included in IntegrAlign pipeline to achieve an optimal alignment, is the evaluation of the precision of the transformation at the single cell level using Napari [16], a multi-dimensional image viewer for python, in full-resolution images. Indeed, a side-by-side display of both images with a duplicated cursor moving over the 2 images (thanks to the transformation of its coordinates) allow the evaluation of alignment at specific coordinates (**Fig. 2.C**), enabling the visualization of the mapping of each coordinate. This approach allows to thoroughly examine the entire slide, assess the overall alignment, and identify any regions where alignment may be problematic.

### 2.4 Image clipping

We also included in this tool the possibility to eliminate regions that differ between the 2 slides (i.e.: tissue tearing or folding, **Supp. Fig. 1**). So, filtering of these regions is a crucial step allowed by IntegrAlign pipeline to get a robust integrated dataset after alignment. Using the previously described side by side visualization with the duplicated cursor, non-alignable tissue can be located and excluded with manual clipping (**Fig. 2D**).

### 2.5 Cells integration

Finally, cell coordinates of one slide are mapped to the other, with exclusion of cells located in non-alignable tissue sections delimited with the manual clipping. This involves transforming cells coordinates from the fixed (reference) image to the moving (target) image. For a straightforward qualitative alignment validation, we can also use a common marker (in our case Cytokeratin) to check if the structure aligns correctly between the slides, ensuring accurate alignment.

### 2.6 Availability

IntegrAlign, developed in Python, is freely available under the MIT license at https://github.com/CAUXlab/IntegrAlign.

## 3 Validation methods

### 3.1 Simulation of a serial slide

In order to evaluate the IntegrAlign accuracy, we artificially generated a corresponding serial slide from the original. To achieve this, we induced random B-spline deformations (mesh size = 3) with maximum control point displacement to simulate potential inter-slide deformations. Then, to replicate varied spatial orientations, specific rotation and translation transformations were also implemented. Using three maximum displacements of the control points (100, 500 and 1000µm) in the B-spline transformation, we obtained three different deformed slides and the corresponding ground truth composite transformation.

### 3.2 Misalignment distance

In order to quantify the quality of the alignment we defined corresponding points in two sl ides and comp uted the distance between trans forme d (from s lide 1) to ground truth coordin ate (slid e 2). L ocal alignment error (µm) was defined a s follows :

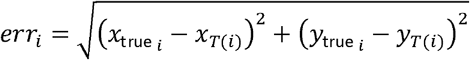

Where *err*_*i*_ represents the misalignment error for the *i*-th point, *x*_*T*(*i*)_and *y*_*T*(*i*)_ are the transformed coordinates and 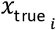 and 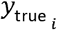 are the ground truth coordinates.

### 3.3 Nearest neighbor method

We computed the distance between every CD8 ^+^ cells and their nearest tumor cell (CK positive) denoted as 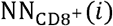:

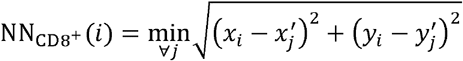

Where (*x*_*i*_, *y*_*i*_) are the coordinates of CD8^+^ cell *I*, (*x*^′^_*i*_, *y*^′^_*i*_) are the coordinates of CK^+^ cell and 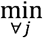 is the minimum over all CK^+^ cells.

### 3.4 Confidence interval on median

To quantify the uncertainty around the median of the distances between CD8 and CK cells, we computed its confidence interval (CI) using the bootstrap method. This approach involved generating 1,000 resampled datasets by randomly selecting nearest neighbor distances (CD8-CK) with replacement from the original distribution. For each resampled dataset, we computed the median. The 95% confidence interval was then determined by calculating the 2.5th and 97.5th percentiles of the medians obtained from these resampled datasets.

### 3.5 Other alignment methods

We compared IntegrAlign efficacy with the following existing methods:

#### Manual

Estimation of a transformation using control point mapping. We utilized the Scikit-image library [17] to estimate a projective transformation based on 100 matching coordinates manually annotated in QuPath [18].

#### Automatic feature detection

To address the limitations of manual annotation, we explored two automatic feature detection algorithms from the OpenCV library [19]: ORB (Oriented FAST and Rotated BRIEF) and SIFT (Scale-Invariant Feature Transform). For ORB descriptors, we employed a brute-force matcher to identify common features while for SIFT descriptors, we utilized a FLANN-based (Fast Library for Approximate Nearest Neighbors) matcher. These methods automatically identify key point descriptors and match the common ones, allowing to estimate the transformation using the RANSAC (Random Sample Consensus) algorithm.

#### Intensity-based optimization method

Another approach to estimate the transformation without manual feature definition is by using intensity-based optimization. For this comparison, we used only the rigid transformation component of IntegrAlign, implemented with SimpleITK [13]. This method corrects for rotation and translation but does not allow for local image deformation.

By comparing these methods with IntegrAlign, we assessed its performance and efficiency, highlighting the differences and potential advantages of our tool in various alignment scenarios.

### 3.6 Mature dendritic cells ground truth coordinates

We established the ground truth coordinates for mature dendritic cells (DCLAMP^+^ cells) by manually annotating the common DC-LAMP positive cells across two serial slides. This annotation process involved checking each coordinate across the entire slide using QuPath [18] and recording indices for cells that match in both slides.

## Results

### 1 On simulated serial slides, the alignment error is smaller than the diameter of the cell nucleus

A natural approach to quantifying the accuracy of IntegrAlign (the alignment error) is to compare the cell coordinates following IntegrAlign alignment to ground truth coordinates. Because ground truth coordinates are generally unknow, this benchmark is impractical. To address this, we start with mIF slide and artificially deformed slide: this allows tracking the coordinates of points in the original and deformed images, so that we can compare slides post-realignment to a ground truth (the applied deformation).

We first evaluate IntegrAlign accuracy using four breast patient slides. For each slide, we simulate serial slides by applying the same composite transformation model used for alignment (**Fig. 4A**). For each initial breast tumor image, we thus generate three different deformed slides and their corresponding ground truth transformations depending on the maximum displacement of the control points (100, 500 and 1000µm) in the B-spline transformation. Deformed slides were then aligned to the original images and the composite transformation estimated using IntegrAlign. To quantify alignment accuracy, we manually defined 100 points in the tissue of the original slide and transformed them in the artificial serial slide (**Fig. 4B**) using the ground truth transformation (artificial) and the transformation from IntegrAlign alignment. From the transformed coordinates, we computed misalignment distances between ground truth points and points post-transformation.

**Figure 4:**
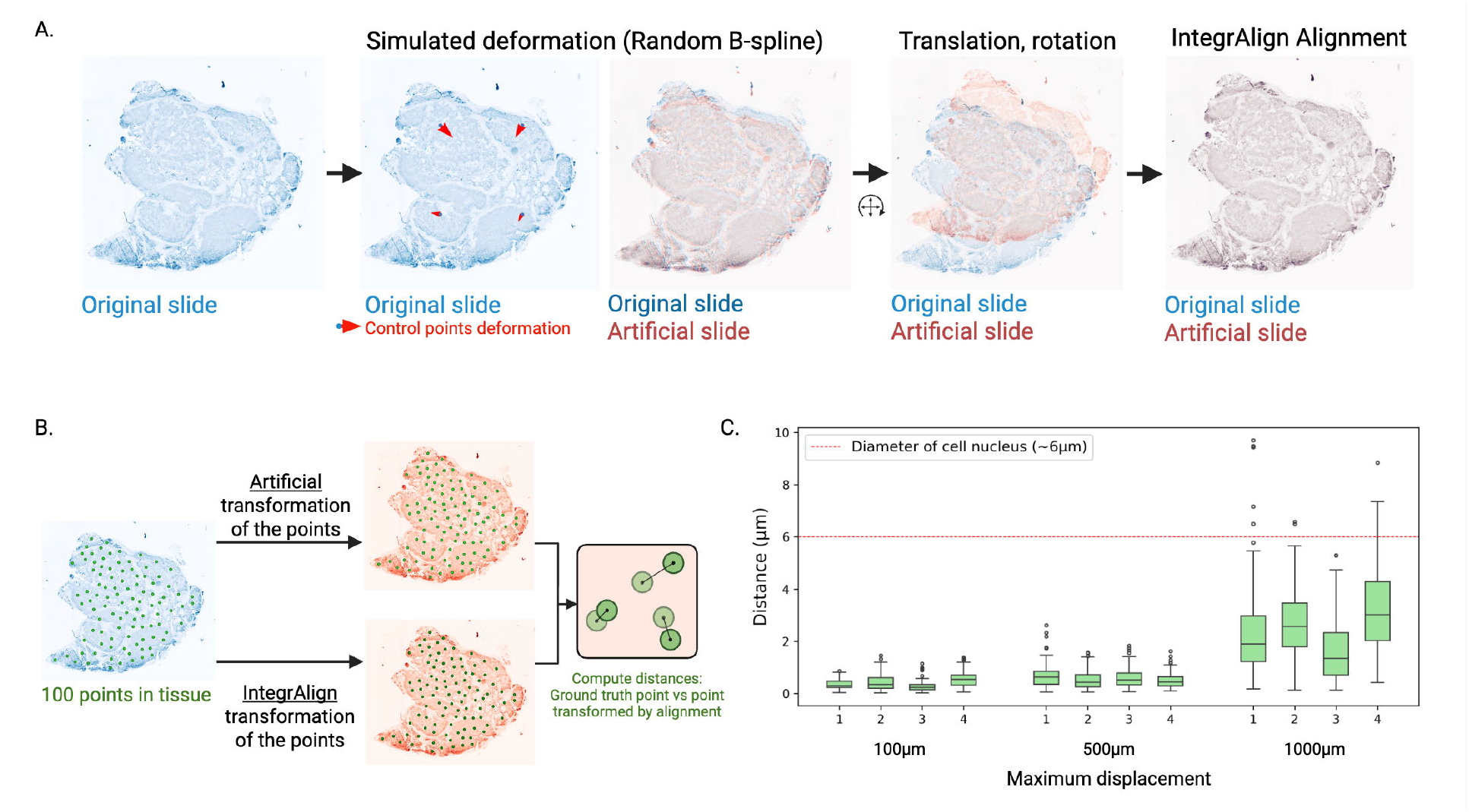
Alignment of an image with an artificially deformed version. **(A)** Random computational deformation of an original slide (blue slide) using B-spline (in this example, a maximum displacement of **1**000µm is shown) and rigid transformations, generating an artificial serial slide (red slide). Then, IntegrAlign alignment is processed between the two synthetic serial slides resulting in the corresponding IntegrAlign transformation **(B)** Evaluation of the IntegrAlign alignment accuracy of the original slide onto the artificial serial slide. A random selection of 100 coordinates in the original slide’s tissue are converted by the artificial transformation or the IntegrAlign transformation into the artificial serial slide. Finally, the distances between transformed coordinates are calculated. **(C)** Box-plots showing the distance between coordinates from the original slide converted with the artificial transformation (from B-spline deformation with 100, 500 or l000µm of maximum displacement; mesh size = 3) and the corresponding coordinates converted with IntegrAlign transformation (coming from alignment). Shown are n=4 representative pairs of synthetic serial slides. Red dotted line represents the mean diameter of cell nuclei as a reference.

**Figure 5:**
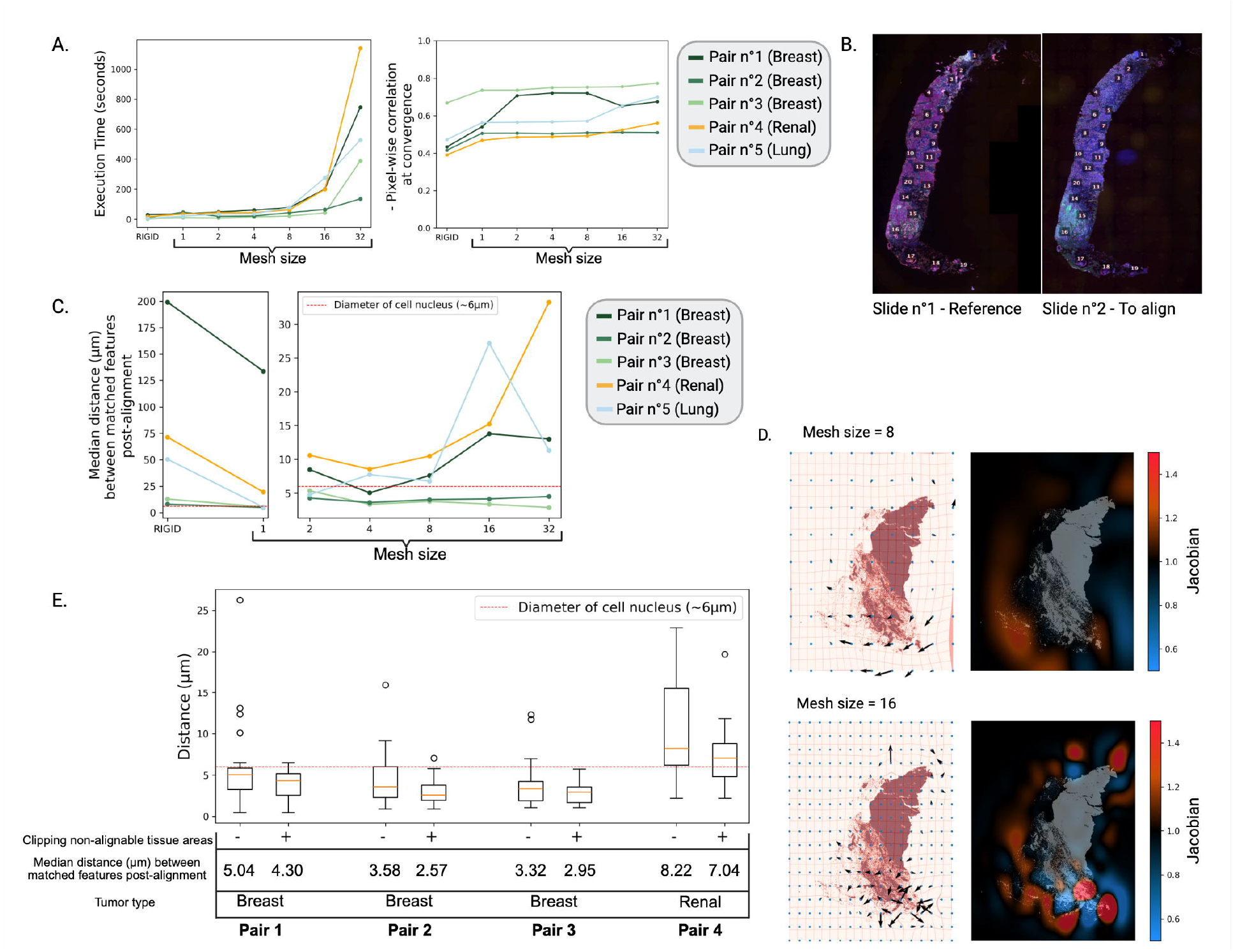
Features and accuracy of IntegrAlign alignment on two serial slides. **(A)** Execution time and pixel-wise correlation of the alignment computed with different mesh sizes on 5 pairs of serial slides from different tissues. Rigid transformation alone is shown as a reference. **(B)** 20 common features are manually marked on slide I (left) and slide n°2 (right) to compute the distances between the coordinates in slide n°2 and the transformed coordinates of slide n°1 in slide n°2. **(C)** Alignment error represented by the median distance (µm) between the common features for different mesh sizes. **(D)** Inverse representation of the position, direction and intensity of deformations induced by the alignment on the resampled target image (deformed target slide) with a deformation grid (left panel) and the map of the jacobian determinant (red = compression, blue = expansion) for a mesh size value of 8 (top panel) or 16 (bottom panel). **(E)** Box-plots showing the alignment error with (+) or without (−) manual exclusion (clipping) of non-alignable tissue areas for each pair of serial slides. The pair 5 is not shown as no non alignable tissue areas were present. Red dotted line represents the mean diameter of cell nuclei as a reference.

Larger deformations reduced alignment accuracy (**Fig. 4C**). Yet, even with large deformations (1000µm displacement), misalignment distance remained low, smaller than the diameter of cell nuclei (6µm). This indicates robust performance of the alignment.

### 2 Alignment accuracy on two serial slides depends on the resolution of deformation and can be optimized by clipping non-alignable tissue areas

We next tested IntegrAlign on authentic serial slides to evaluate its performance on real-world data.

Five pair of serial slides were aligned using only Rigid transformations (step 1) or combining Rigid and B-spline transformation (steps 1 & 2, IntegrAlign pipeline). For the B-spline transformation, we varied the mesh size parameter from 1 to 32, allowing for different levels of deformation complexity (a higher mesh size allows more complex deformations).

Augmenting the mesh size increased execution time exponentially (**Fig. 5A – left**) while pixel-wise correlation (inverse of the similarity metric at convergence) tended to level-off (**Fig. 5A – right**). To quantify the alignment quality, we manually annotated twenty common features between each pair of serial slides using QuPath [18], as illustrated in **Fig. 5B**. These common features provided us with corresponding ground truth coordinates for each slide, allowing the computation of the alignment error (**Fig. 5.C**). Increasing the mesh size from 1 to 8 initially increased accuracy, but with higher mesh sizes deteriorated alignment precision in a slide-specific manner (**Fig. 5.C**). As shown in the deformation grid and by quantifying expansions and compressions using the determinant of the Jacobian (Methods), a mesh size of 8 induced coherent deformations, while a higher mesh size infers erratic deformations inside the tissue (**Fig. 5D, Supp. Fig. 1.A**).

Examining alignment at single-cell resolution by displaying serial slides side by side (Napari dual viewer, **Fig. 2.C**) and mirroring the cursor of the fixed slide into the moving slide showed alignment was often problematic in regions with folds or tearing present in one slide but not in the other (see examples in **Supp. Fig. 1**). Clipping of these zones resulted in a diminution of the overall alignment error (**Fig. 5E**).

### 3 The precision alignment provided by IntegrAlign opens the door to combined spatial analyses, including nearest neighbor analysis

The ultimate goal of integrating serial slides employing multiple mIF panels is to perform combined spatial analysis on extended cell types and phenotypes. To evaluate the feasibility of spatial analyses combining serial slides, we performed nearest neighbor analysis on a pair of serial slides (breast tissue). We computed the distances between CD8^+^ T cells and closest tumor cells (CK positive) from slide 1 (CD8.1 / CK.1) and slide 2 (CD8.2 / CK.2) within each slide, to serve as ground truth. We then performed the same analysis between CD8^+^ T cells from the slide 1, transformed to the coordinate system of slide 2, and tumor cells of slide 2 (CD8.TR1-2 / CK.2), combining them into a single coordinate system (**Fig. 6A-B**). Nearest neighbor distances distributions were similar across these two scenarios (**Fig. 6C**). For example, median CD8-tumor cell distances were 23.1 µm (IC: [21.5, 24.4]) and 22.4 µm (IC: [21.4, 23.3]) within slide 1 and slide 2 respectively, compared to 22.5 µm (IC: [21.4, 23.8]) across aligned slides. The median absolute difference between distances of CD8.1 / CK.1 and CD8.TR1-2 / CK.2 was 4.6 µm (**Fig 6D, Supp. Fig. 2**), an acceptable error relative to the median CD8-tumor cell distance (23.1 µm).

**Figure 6:**
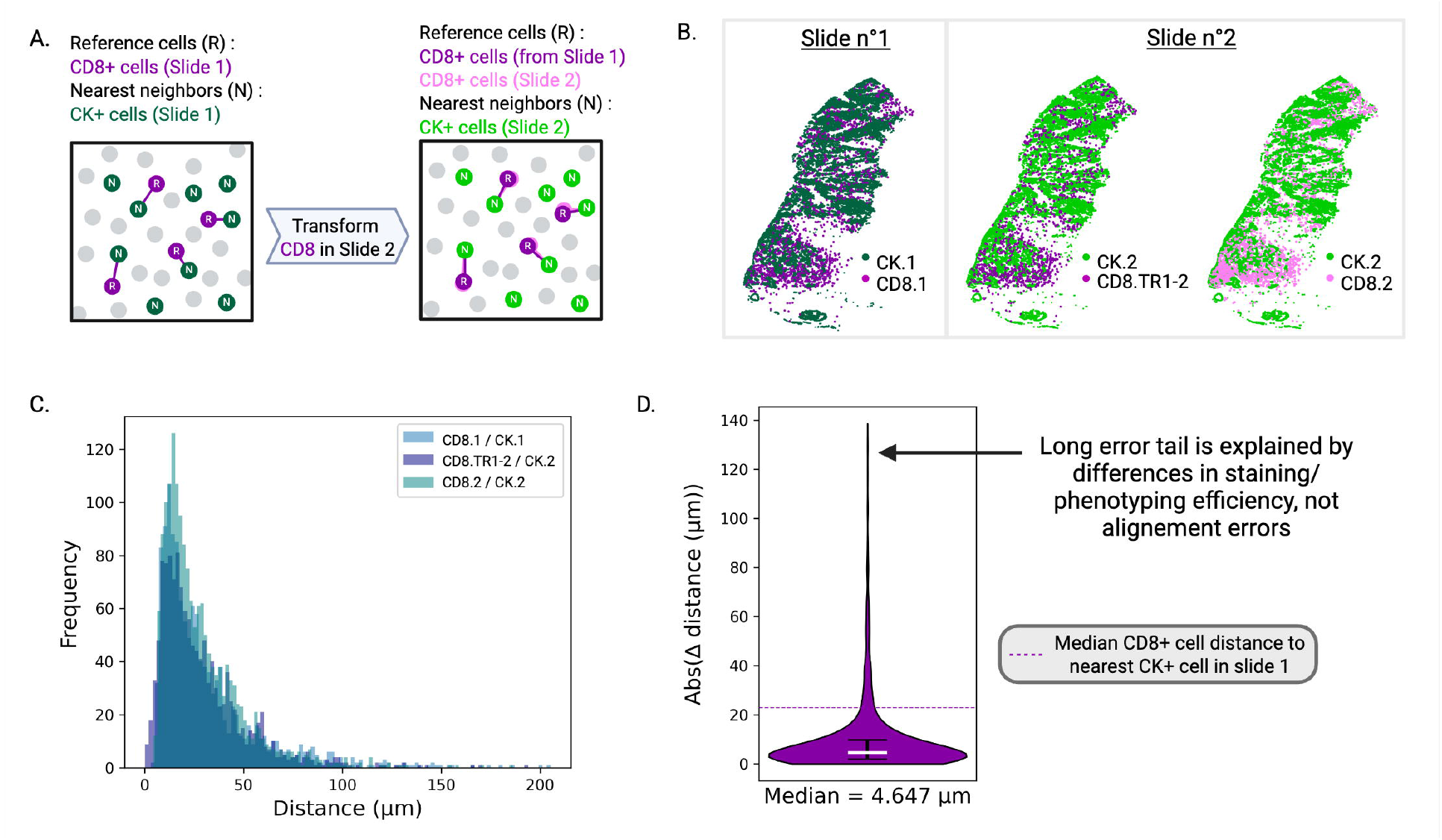
Nearest neighbor analysis of CDS+ T cells and tumor cells (common markers) between two serial slides aligned by IntegrAlign. **(A)** Schematic illustration of the nearest neighbor distances of reference CD8+ cells to neighbor tumor (CK +) cells between two serial slides. The analysis includes computation of the distances between CD8+ and nearest CK+ cells within a single slide (n°1 or n°2) and also between CD8+ cells from slide n°1 transformed in slide n°2 and nearest CK+ cells of slide n°2. **(B)** Visual representation of the spatial distribution of cell coordinates in both slides. CK.1, CD8.1 are cells from slide n°1. CK.2, CD8.2 are cells from slide n°2. CD8.TR1-2 are the transformed coordinates of CD8.1 in the coordinate system of slide n°2. **(C)** Distribution of the nearest neighbor distances for each CD8-CK pair visualized in **(B)**. **(D)** Absolute value of the distance differential (absolute error) between CD8 of slide n°1 to the nearest CK of slide n°1 and CD8 from slide n°1 in slide n°2 to the nearest CK of slide n°2.

### 4 Unprecedented alignment accuracy using IntegrAlign compared to existing alignment methods

To benchmark IntegrAlign together with other alignment methods, we used two serial slides from one breast cancer patient and compared alignment based on (i) manual annotation of common features, (ii) ORB and SIFT [19], two automatic features detection algorithms, (iii) Rigid transformation (IntegrAlign alignment - step 1 only) and (iv) the complete IntegrAlign pipeline (Rigid + B-spline transformations).

Ground truth coordinates were defined by manually annotating the DC-LAMP^+^ cells (corresponding to mature dendritic cells) common to both serial slides (**Fig. 7A**). By transforming DC-LAMP positive cells coordinates from slide n°1 to slide n°2 using each estimated transformation (**Fig. 7A**) and computing every corresponding misalignment distance with the ground truth cell coordinates in slide n°2, we find that IntegrAlign most accurately aligns serial slides: alignment error was 7µm with IntegrAlign compared to 38-200 µm with other approaches. Much of this gain seems to be attributable to step 2 of IntegrAlign (local deformation with B-spline transformation): the entire IntegrAlign pipeline reduced the misalignment distance by 92% compared to the Rigid transformation only (step 1 of IntegrAlign; Rigid-only error: 89 µm, entire IntegrAlign pipeline: 7 µm).

**Figure 7:**
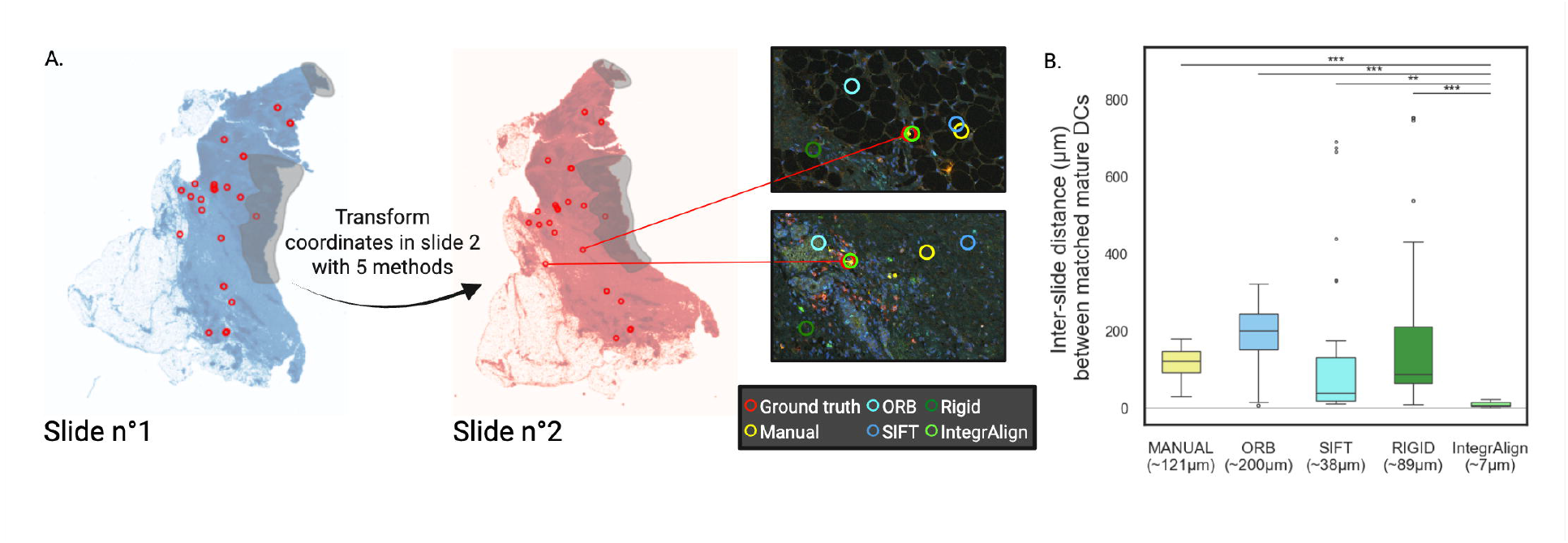
Comparative analysis of alignment methods. Comparison of lntegrAlign with other registration methods performed on one pair of serial slides. **(A)** Manual identification of shared mature dendritic cells (red circles) between two serial slides. Coordinates of slide n°1 are transformed into slide n°2 using five different methods: Manual (projective transformation of common features annotated manually), **ORB** and SIFT (projective transformation of common features annotated automatically), Rigid (alignment of images based on their pixel instensity with optimization algorithm using translation and rotation), IntegrAlign (alignment of images based on their pixel instensity with optimization algorithm using translation, rotation and deformation). **(B)** Box-plots showing the distances between the real dendritic cell coordinates of slide n°2 compared to the transformed coordinates from slide n°1 into slide n°2’s coordinate system using the different methods. Welch’s t-test indicates that lntegrAlign significantly reduces misalignment distance compared to the Manual method (p = 8.328e-15), **ORB** method (p = 1.602e-11), SIFT method (p = 3.779e-03), and Rigid method (p = 1.602e-11).

## Discussion

Assessing the accuracy of alignment methods is inherently challenging due to the lack of reliable common descriptors between serial slides. Indeed, even if they pertain to one tissue biopsy, the separation between serial slides (4µm) results in differences at the cellular level, making it impossible to perform direct comparison by retrieving all cells from one slide and contrasting them to the other slide. To overcome this, one can manually define common features between slides. In **Fig. 7**, we achieved this by manually matching authentic cell coordinates, which is arguably the most representative method we used, and IntegrAlign demonstrated promising results using this approach. Indeed, compared to existing methods that use only rotation and translation, IntegrAlign also incorporates local deformation with the sequential application of Rigid and B-spline transformations from the Simple ITK package. By adopting this two-step strategy, we effectively reduced the risk of unwanted random deformations that may arise when directly applying the B-spline transformation to severely misaligned images. This approach ensures a smoother optimization process and enhances the overall alignment outcome, providing a more robust and accurate alignment of the images. Also, regarding the alignment itself, compared to the commonly used methods that need annotation of matching control points to estimate the transformation, which is time-consuming and labor-intensive particularly when high precision is required, IntegrAlign alignment is fully automated thanks to the use of the intensity-based optimization algorithm.

In developing IntegrAlign, we made methodological decision to optimize computational efficiency, which make IntegrAlign suitable to process imaging data from large cohorts. We chose the L-BFGS-B optimizer which efficiently approximates the inverse Hessian matrix, without explicitly computing or storing it. This is beneficial since we are dealing with large-scale optimization problems where memory usage is a concern. This keeps memory requirements manageable while speeding up convergence compared to first-order methods like gradient descent. Among the various similarity metrics that the SimpleITK [13] package can use to optimize B-spline deformation, we found that correlation optimized computation time while performing well in terms of accuracy (*data not shown*).

Computing this similarity metric on only a fraction of random sampled pixels combined with transient downscaling the imaging data (DAPI channel) significantly decreased computation time and memory while preserving the information needed for accurate alignment. Larger mesh size can lead to an exponential increase of the execution time (**Fig. 5A – left**), an issue when aligning a large number of samples. Larger mesh sizes also endow slide alignment with too much freedom, resulting in «over-alignment” where tissue stretching occurs to match areas that do not correspond because it increases the similarity metric (**Fig. 5D**). Such discrepancies can arise due to slight technical differences between scanning of serial slides, random staining variations, or differences in tissue content (damaged tissue) between slides. Therefore, mesh sizes ranging from 1 to 10 can optimize computational efficiency without trade-off in terms of accuracy.

### Key considerations

In the context of application to large cohorts of samples, it is important to identify specific cases and slides that cannot be aligned beforehand to ensure proper alignment and avoid wasting time.

Also, when dealing with images containing multiple pieces of tissue, alignment often fail due to varying distances between these pieces. A practical solution would be to crop the image to isolate each tissue piece and align them one by one. This ensures more accurate alignment by addressing each tissue section’s specific features independently.

As shown in **Fig. 1A** the final step of IntegrAlign involves transforming the cell coordinates from the fixed (reference) image to the moving (target) image. Alternatively, we can specifically transform cells coordinates from the moving to the fixed image, but this requires computing the inverse transformations. Therefore, choosing which slide serves as the fixed and moving image is crucial as it determines the target image where all coordinates will be transformed. The target image should be the most important panel as it will not have any alignment error since the coordinates are not transformed.

### Opportunities for Improvement

Slides with irregular tissue tearing across the entire slide may not align correctly and should be removed before proceeding. The decision can be made by visualizing the downscaled DAPI images to assess the degree of variation between serial slides. However, if the tearing is regular compared to the other serial slides and affects only a portion of the slide or if there is identified staining issues that can lead to certain areas not being analyzed (manual annotation of artefacts), we can address these issues with the annotations of empty regions and artefacts. By integrating the annotations into the coordinate system of the target image, we can retain only the cells located in the unblemished tissue of both slides. Indeed, the combined annotation map will accurately represent the tissue structure ensuring that cells present in one slide but transformed into empty areas or artefact in another can be properly identified and managed. Enabling the removal of cells that are erroneously placed in regions with artefacts or missing tissue will automate image clipping and enhance the accuracy of subsequent analysis and interpretation.

In order to choose the optimal mesh size, we verify that it does not induce too much deformation with the deformation grid and Jacobian analysis. Additionally, we could examine the correlation factor between overlaying Rasters of cell coordinates (from DAPI) from both slides. While the manual annotation of common features is highly time-consuming and labor-intensive, using the correlation factor would be a promising automatic approach for assessing alignment accuracy across the entire slide.

We could extend the alignment process beyond two slides to include three serial slides and increase even more the potential number of identified cell types and therefore the possibilities in the subsequent spatial analyses. It would allow to capture finer details and variations in tissue structure and cellular composition.

Additionally, by using this method, we will approach the number of parameters seen in advanced techniques like CODEX and IMC [20], which focus on specific regions of interest (ROIs). However, with augmented multiplex immunofluorescence, we analyze whole slides, providing a more comprehensive representation of the spatial architecture. This approach is particularly beneficial in oncology for example, where large cohorts of patients are studied and direct analysis of the entire slides avoid bias from selecting specific regions. The technique is cost-effective, less time consuming and can be translated to clinical routine in particular to analyze immune infiltrates and biomarkers in whole slides.

With an augmented number of markers, more complex spatial analyses become possible such as the identification of niches with tools like NIPMAP: niche-phenotype mapping of multiplex histology data by community ecology [21].

