## Supplementary figures and images for "IntegrAlign: A comprehensive tool for multi-immunofluorescence panel integration through image alignment"

### Supp. Fig. 1

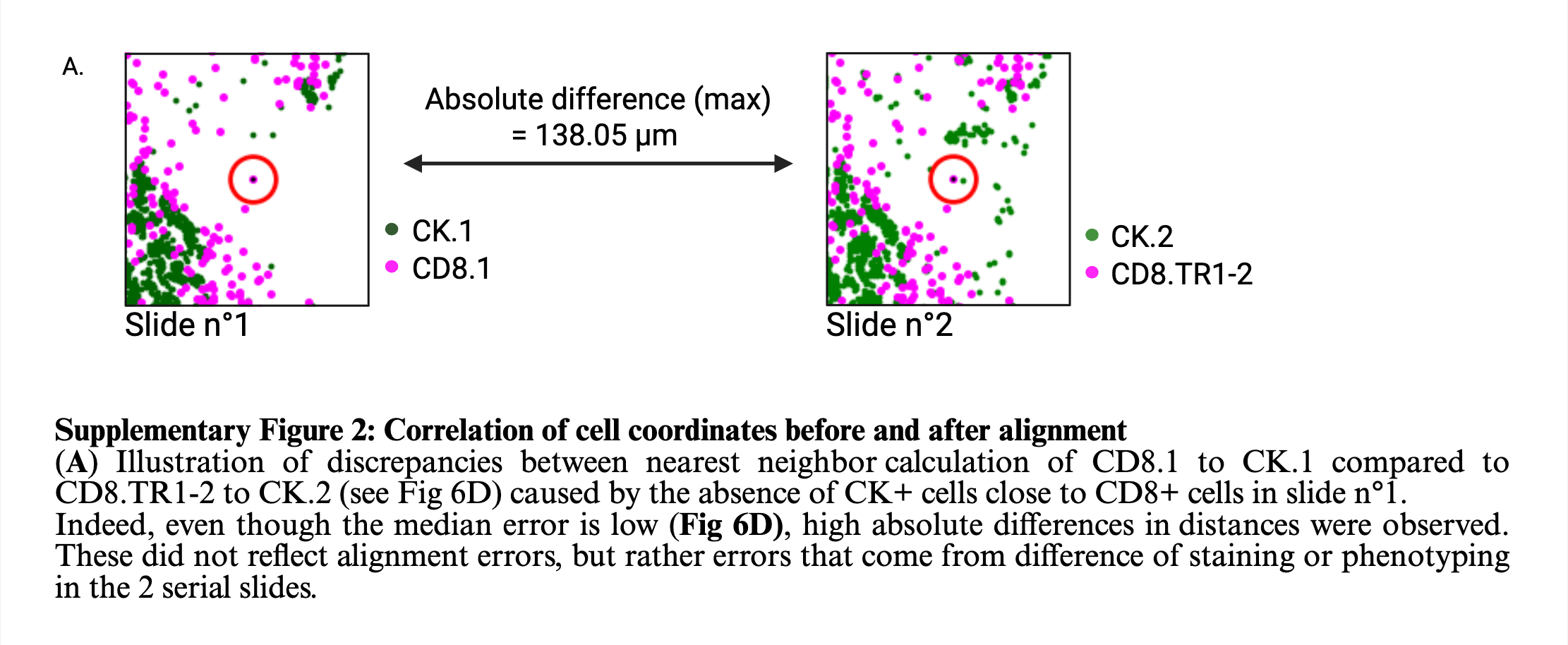

### Supp. Fig. 2

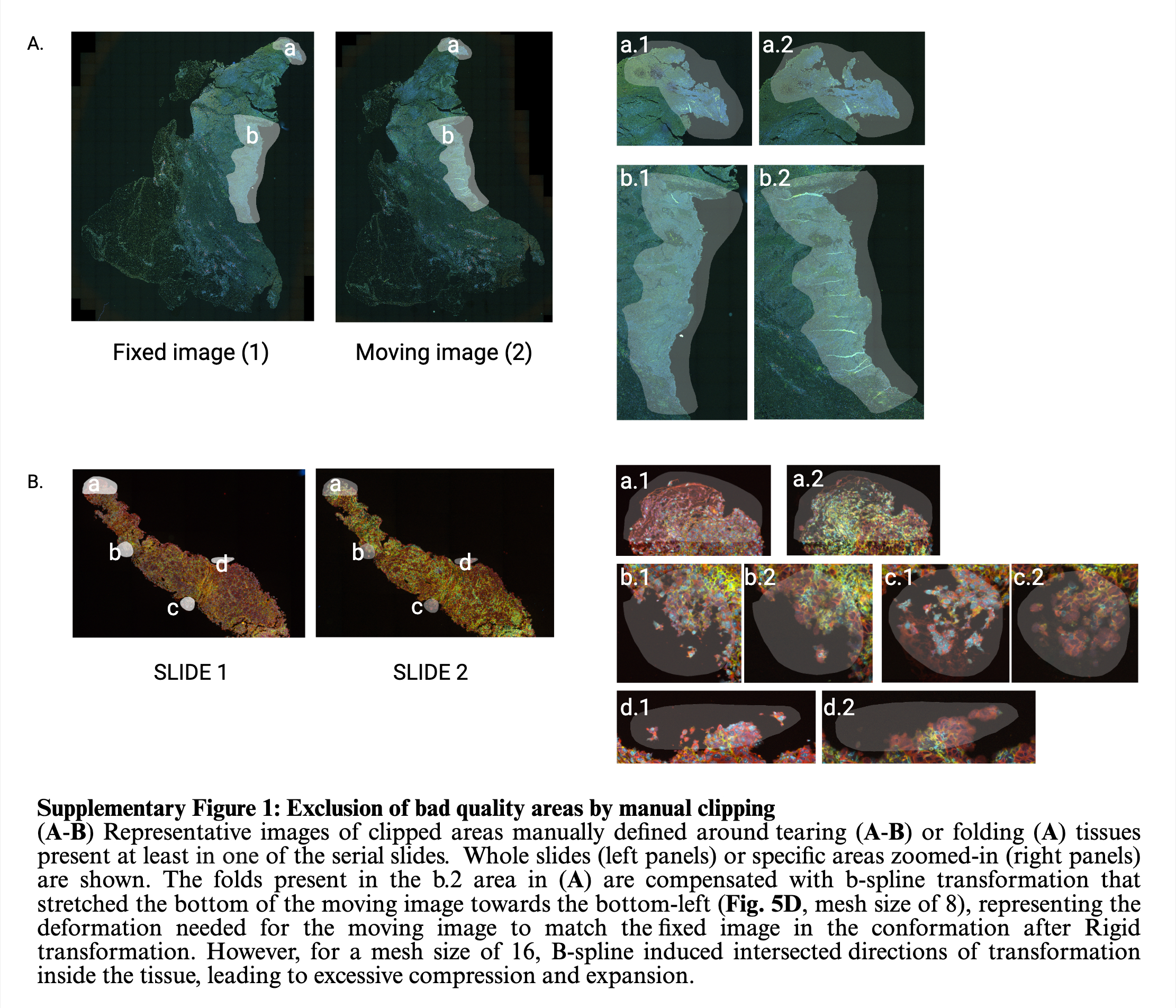
